# Driving factors of conifer regeneration dynamics in eastern Canadian boreal old-growth forests

**DOI:** 10.1101/2020.02.26.966200

**Authors:** Maxence Martin, Miguel Montoro Girona, Hubert Morin

**Author notes:** **Author Contributions:** Conceptualization: MM, MMG, HM. Fieldwork: MM, MMG. Data curation: MM. Investigation: MM. Methodology: MM. Data analyses: MM. Results interpretation: MM. Project administration: MM. Resources: HM. Supervision: HM, MMG. Visualization and edition: MM. Writing–original draft: MM, MMG, HM. Writing–review: MM. Funding: HM, MMG. **Funding:** Funding was provided by the Natural Sciences and Engineering Research Council (NSERC) of Canada, the Canada Research Industrial Chairs Program obtained by HM, “Chaire de recherche industrielle du CRSNG sur la croissance de l’épinette noire et l’influence de la tordeuse des bourgeons de l’épinette sur la variabilité des paysages en zone boréale,” as well as a new professor start-up fund at University of Québec in Abitibi Temiscamingue (UQAT) and a silvicultural research grant from MRC-Abitibi that were both obtained by MMG.

## Abstract

Old-growth forests play a major role in conserving biodiversity, protecting water resources, sequestrating carbon, and these forests are indispensable resources for indigenous societies. To preserve the ecosystem services provided by these boreal ecosystems, it becomes necessary to develop novel silvicultural practices capable of emulating the natural dynamics and structural attributes of old-growth forests. The success of these forest management strategies depends on developing an accurate understanding of natural regeneration dynamics. Our goal was therefore to identify the main patterns and the drivers involved in the regeneration dynamics of old-growth forests, placing our focus on boreal stands dominated by black spruce (*Picea mariana* (L.) Mill.) and balsam fir (*Balsam fir* (L.) Mill.) in eastern Canada. We sampled 71 stands in a 2200 km^2^ study area located within Quebec’s boreal region. For each stand, we noted tree regeneration (seedlings and saplings), structural attributes (diameter distribution, deadwood volume, etc.), and abiotic (topography and soil) factors. We observed that secondary disturbance regimes and topographic constraints were the main drivers of balsam fir and black spruce regeneration. Furthermore, the regeneration dynamics of black spruce appeared more complex than those of balsam fir. We observed distinct phases of seedling production first developing within the understory, then seedling growth when gaps opened in the canopy, followed by progressive canopy closure. Seedling density, rather than the sapling density, had a major role in explaining the ability of black spruce to fill the canopy following a secondary disturbance. The density of balsam fir seedlings and saplings was also linked to the abundance of balsam fir trees at the stand level. This research helps explain the complexity of old-growth forest dynamics where many ecological factors interact at multiple temporal and spatial scales. This study also improves our understanding of ecological processes within native old-growth forests and identifies the key factors to consider when ensuring the sustainable management of old-growth boreal stands.

## Introduction

The global extent of native old-growth forest has declined markedly over the past few centuries through a cumulative and increasing impact from anthropic activities within these forest landscapes (1–3). The boreal forest, most of which is situated in Canada and Russia, is currently the largest reserve of natural forest on our planet (3). Boreal old-growth forest has also experienced rapid loss over the last centuries (1,4,5). The remaining old-growth forests are critically important to biodiversity, water resources, carbon sequestration and storage, and these stands remain integral elements of indigenous societies and even human health (3, 6). The sustainable management of boreal forests has a primary goal of protecting the remaining old-growth forests. Restoring the integrity of intact forests is also an urgent issue; this is especially true in Fennoscandia where old-growth forests have been almost completely eliminated (7). We are therefore facing a critical situation where novel silvicultural practices and restoration strategies are now priorities in the context of the global biodiversity crisis, climate change, and forest sustainability.

Effective forest restoration strategies require an accurate understanding of the natural dynamics of old-growth forests. Tree regeneration is an essential process in forest ecosystems to ensure the persistence and resilience of forest stands when subjected to various disturbances (8, 9). As such, forest science is placing increased importance on understanding tree regeneration following natural and anthropic disturbance (e.g., 10–16). However, regeneration dynamics in old-growth forests remain an understudied subject in ecology; this absence is particularly true for the boreal biome. Moreover, due to the scarcity of old-growth stands in many boreal regions, conducting studies related to this subject is often challenging, given the lack of reference sites. This need for baseline data underscores the important scientific value of the boreal biome in eastern Canada where some regions still contain large intact stands of forest as intensive forest management practices only began relatively recently, i.e., since the 1960s (17, 18). The study of regeneration dynamics in the boreal old-growth forests of eastern Canada thus represents a benefit for all boreal regions, especially those where these ecosystems have been almost completely eliminated. Black spruce (*Picea mariana* (L.) Mill.) and balsam fir (*Abies balsamea* (L.) Mill.) are the two main late-successional species in the eastern Canadian boreal forest (19). Pure black spruce or mixed black spruce–balsam fir stands are the most common old-growth forest types in eastern Canada (19–21). Old-growth forests are also, however, the most logged forest type in this territory, leading to the rapid loss of old-growth forest surfaces (5,22,23). Pure black spruce stands are under even greater pressure as this specific old-growth forest type is most selected for logging given the high economic value of this species (23).

Both black spruce and balsam fir are well adapted to long (>150 years) periods of suppressed growth in the understory (24–26). These species are also able to regenerate under their own cover, mostly through vegetative reproduction for black spruce—regeneration by layers—and sexual reproduction, i.e., seed origin, for balsam fir (19). Previous studies have highlighted that the seedling densities of black spruce and balsam fir are similar under gaps or canopy cover (27–29). When a gap in the canopy opens as a result of a secondary disturbance, the gap-fillers will therefore generally be pre-established regeneration rather than seeds or layers that would have established following the disturbance (30, 31). Once a gap is created, the regeneration trees of both species increase their vertical growth to reach the overstory relatively quickly (26,32,33). However, black spruce and balsam fir differ in their ecological strategies in terms of growth, sensitivity to disturbance, resistance to fire, and seed dispersal; as such, these differences should vary their specific regeneration dynamics. Balsam fir regeneration is seen as being more competitive than that of black spruce due to balsam fir seedlings’ faster and more intense growth response to canopy openings (31, 34). Balsam fir, however, is more vulnerable to spruce budworm (*Choristoneura fumiferana* (Mills.)) outbreaks, windthrow, and root rot than black spruce (35–38). Moreover, balsam fir seeds are not adapted to fire, making this species strongly dependent on the proximity of seed trees, as opposed to black spruce that is very well adapted to fire events (39). Black spruce also outcompetes balsam fir on wet soils (39).

From the abovementioned observations, stands in the old-growth forests in eastern Canada are expected to shift between black spruce–dominated stands and black spruce–balsam fir mixed stands over time (21,28,40). As well, the structure of these stands varies over time (decades and centuries), even though tree species’ composition remains the same (40, 41). At a decennial scale, it is therefore likely that the characteristics of the understory, e.g., tree density or tree species composition within the regeneration layer, will change significantly and rapidly due to the succession of tree-mortality and canopy closure phases.

Understanding the regeneration process in old-growth forests is therefore critical for developing management strategies and silviculture treatments that limit differences between managed and unmanaged forests (42). Our study objective is to identify the main patterns and factors involved in the regeneration dynamics of black spruce and balsam fir in the eastern Canadian boreal old-growth forests. We hypothesize that (1) for both black spruce and balsam fir, sapling density will increase in relation to the secondary disturbance–related structural changes, such as an opening of the canopy and an increase in deadwood volume, and (2) the main differences between black spruce and balsam fir regeneration dynamics are due to abiotic constraints and the availability of proximal balsam fir seed trees.

## Materials and methods

### Study area

Our study involved a 2200 km^2^ region of public forest southeast of Lake Mistassini, Québec, Canada, (**Fig 1**) within an area extending between 50°07ʹ23ʺN to 50°30ʹ00ʺN and 72°15ʹ00ʺW to 72°30ʹ00ʺW. The study zone is crossed by the Mistassini, Ouasiemsca, and Nestaocano rivers and lies within the western subdomain of the black spruce–feather moss bioclimatic domain (43). Regional climate is subarctic with a short growing season (120–155 days). Mean annual temperature ranges between −2.5 and 0.0 °C, and mean annual precipitation is 700 to 1000 mm (43). Surficial deposits consist mainly of thick glacial tills, forming a low-lying topography characterized by gentle hills that vary in altitude from 350 to 750 m asl (44). Black spruce and balsam fir dominate the stands across this territory, while jack pine (*Pinus banksiana* Lamb.), white spruce (*Picea glauca* (Moench) Voss), paper birch (*Betula papyrifera* Marsh.), and trembling aspen (*Populus tremuloides* Michx.) are the secondary tree species.

**Fig 1.**
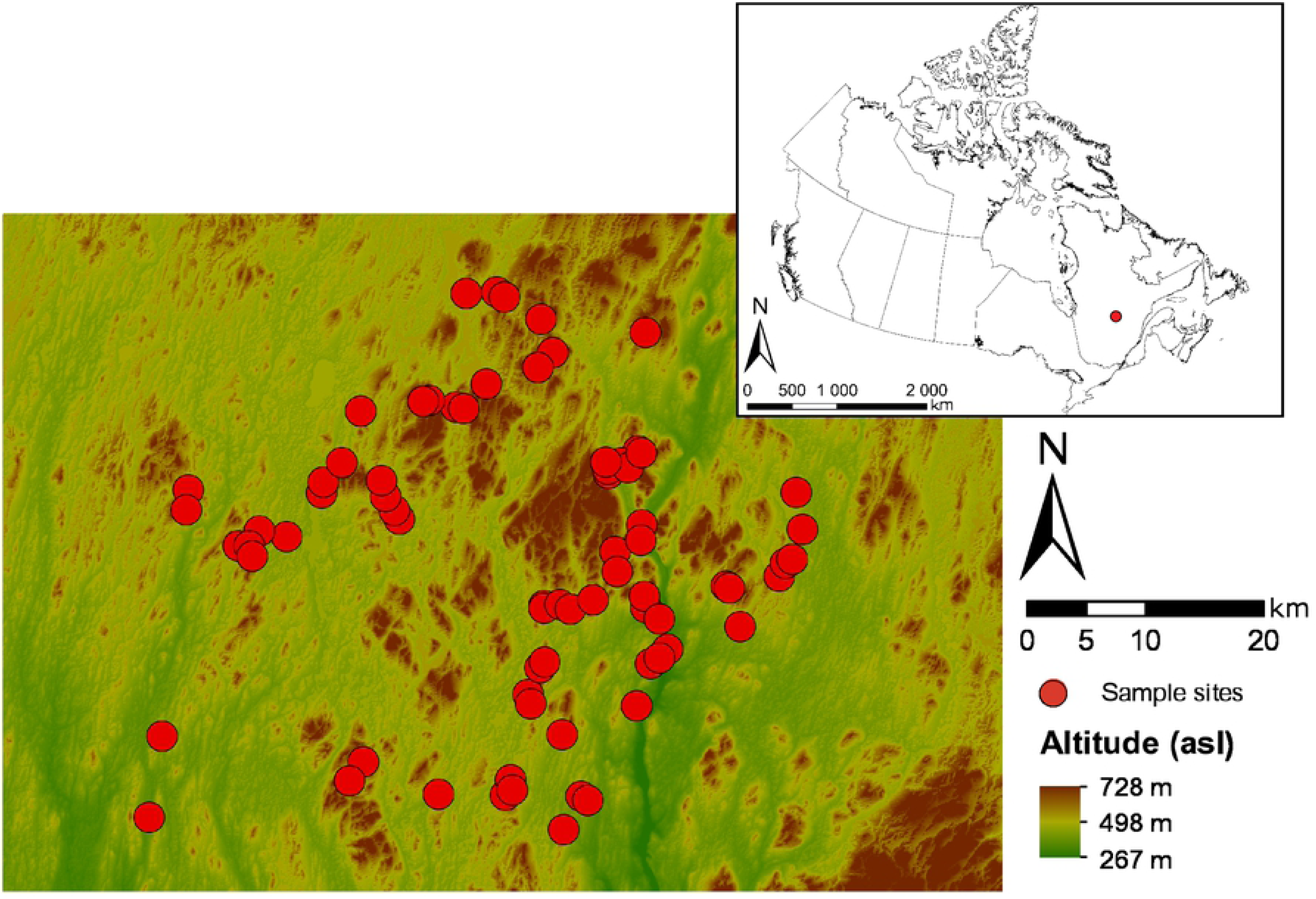
Map of the study territory showing the location of the sample sites (red filled circles). The insert map indicates the location of the study territory in Quebec, Canada (red circle).

Fire is the main driver of stand-replacing disturbances on this territory (45), while spruce budworm outbreaks are the principal agent of secondary disturbance (26). This territory was unmanaged until 1991 when intensive timber exploitation began. The surface area harvested remained relatively low until 2000; however, harvesting increased significantly after this date.

### Experimental design

We sampled 71 stands in the study area during 2015 and 2016 and applied a stratified random sampling approach. Site selection considered two main criteria: 1) that sites reflected the six dominant environmental types found within the study area, according to the ecological classification of the Quebec Ministry of Forests, Wildlife and Parks (MFWP) (43), and 2) that sites must contain two minimal stand-age classes (80–200 years and >200 years). Environmental types are defined through a combination of site potential vegetation, slope classes, surface deposits, and drainage classes. The six dominant MFWP environmental types covered more than 72% of the productive forest. They included: 1) balsam fir–white birch potential vegetation having moderate slopes, till deposits, and mesic drainage; 2) black spruce–balsam fir potential vegetation having moderate slopes, till deposits, and mesic drainage; 3) black spruce–feather moss potential vegetation (BSFM) having gentle slopes, sand deposits, and xeric drainage; 4) BSFM having gentle slopes, till deposits, and mesic drainage; 5) BSFM having gentle slopes, till deposits, and subhydric drainage; and 6) BSFM having gentle slopes, organic deposits, and hydric drainage.

The age classes correspond to the successional stages of the transition process toward the old-growth stage in Quebec boreal forests (20,46,47): 80–200 years (beginning of the transition toward an old-growth forest) and >200 years (end of the transition to an old-growth forest). Stand age was assessed by surveys in 2015 and 2016, during which we collected cores from the root collar of five dominant or codominant trees per site. Tree age was determined from tree-ring counts of these cores using a binocular microscope.

As the study area is very remote and has limited road access, we added additional logistical criteria to the site selection process; we therefore sampled only sites that were accessible via the existing road network. As well, our surveys were systematically placed at 125 m from the stand edge to limit the influence of the edge effect.

### Plot measurements

At each site, we established a permanent square plot (400 m^2^) as the basis for all subsequent transects and subplots (**Fig 2**). We sampled all merchantable trees—trees having a diameter at breast height (DBH) ≥9 cm—in each 400-m^2^ plot. The attributes sampled were species, DBH and vitality (alive or dead). We then surveyed all saplings—stems having a DBH <9 cm and height ≥1.3 m—within two 100-m^2^ (10 m × 10 m) subplots within the larger plot (**Fig 2**). The attributes sampled for saplings were species and DBH. To count seedlings and quantify their attributes, we established twenty-five 4-m^2^ circular plots along five 25-m-long transects (5 circular plots/transect) that extended out from the center of the 400-m^2^ plot. The angle between two neighboring 25 m-long transects was equal to 72°. Transect 1 was the transect oriented due north. Along a transect, the first circular plot was placed 5 m from the center of the 400-m^2^ plot, with the following circular plots separated by 5 m. In each 4m^2^ plot, we inventoried all seedlings by tree species. We also measured gap length along the five 25 m-long seedling transects. We defined the size of our study from other similar studies and the forest survey methods of the Quebec provincial government (15, 48).

**Fig 2.**
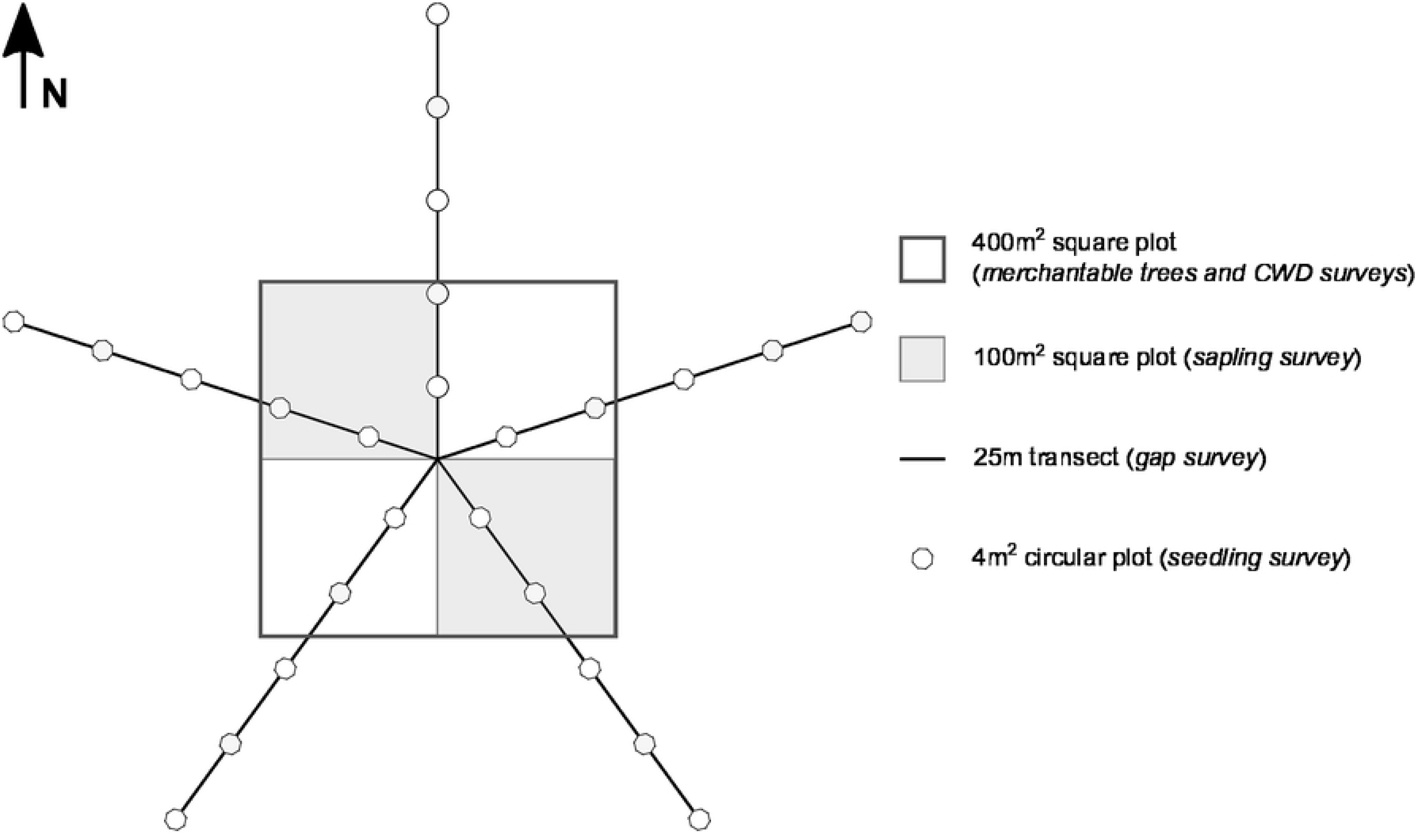
Schematic representation of the experimental design used for the sample sites. N: north; CWD: coarse woody debris.

In addition to these sapling transects, we surveyed coarse woody debris along four 20 m-long transects that followed the edges of the 400-m^2^ plot. We surveyed the diameter of any coarse woody debris intersecting with the transect. We recorded this information for only debris having a diameter >9 cm at the transect intersection. Debris items buried in the organic layer at a depth >15 cm were not sampled. We determined the soil and topographic parameters by digging a soil profile at the center of the 400-m^2^ plot. We used a clinometer to measure slope.

### Data compilation

We applied the following equation to estimate regeneration attributes, i.e., seedling and sapling density, for black spruce and balsam fir:

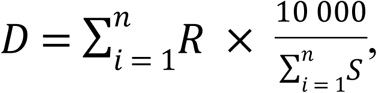

where *D* corresponds to the density per hectare, *R* is the number of seedlings or saplings sampled in each of the *n* plots surveyed, and *S* represents the surface (in m^2^) of each of the *n* plots. (40) had previously computed several structural and environmental attributes for each of the sampled sites used in this study (**Table 1**). Some of these attributes relate to stand structure, including merchantable tree density, basal area, Weibull’s shape parameter of diameter distribution (49), and gap fraction, i.e., the ratio between gap length and total transect length, sensu (50). Other attributes relate to stand composition, such as the basal area proportion of balsam fir. For estimating deadwood, (40) computed the volume of coarse woody debris per hectare using the formula of Marshall et al. (51); however for this study, we also calculated the basal area of snags, i.e., merchantable dead trees, at each study site, an attribute absent from the earlier (40) study. We evaluated forest succession from the minimum time since the last fire, i.e., the age of the oldest tree sampled, and the cohort basal area proportion (CBAP; sensu 52). The latter attribute is an indicator of the stand transition from an even-aged to old-growth stage, i.e., the stage where almost all trees of the first cohort following the last stand-replacing disturbance have disappeared. A CBAP ≈ 0 indicates a stand where all trees belong to the first cohort, and a CBAP = 1 indicates a stand where the first cohort has been entirely replaced by a new shade-tolerant cohort. Finally, we detailed the topographic and pedologic characteristics of the studied stands using slope and the depth of the organic horizon.

**Table 1.**
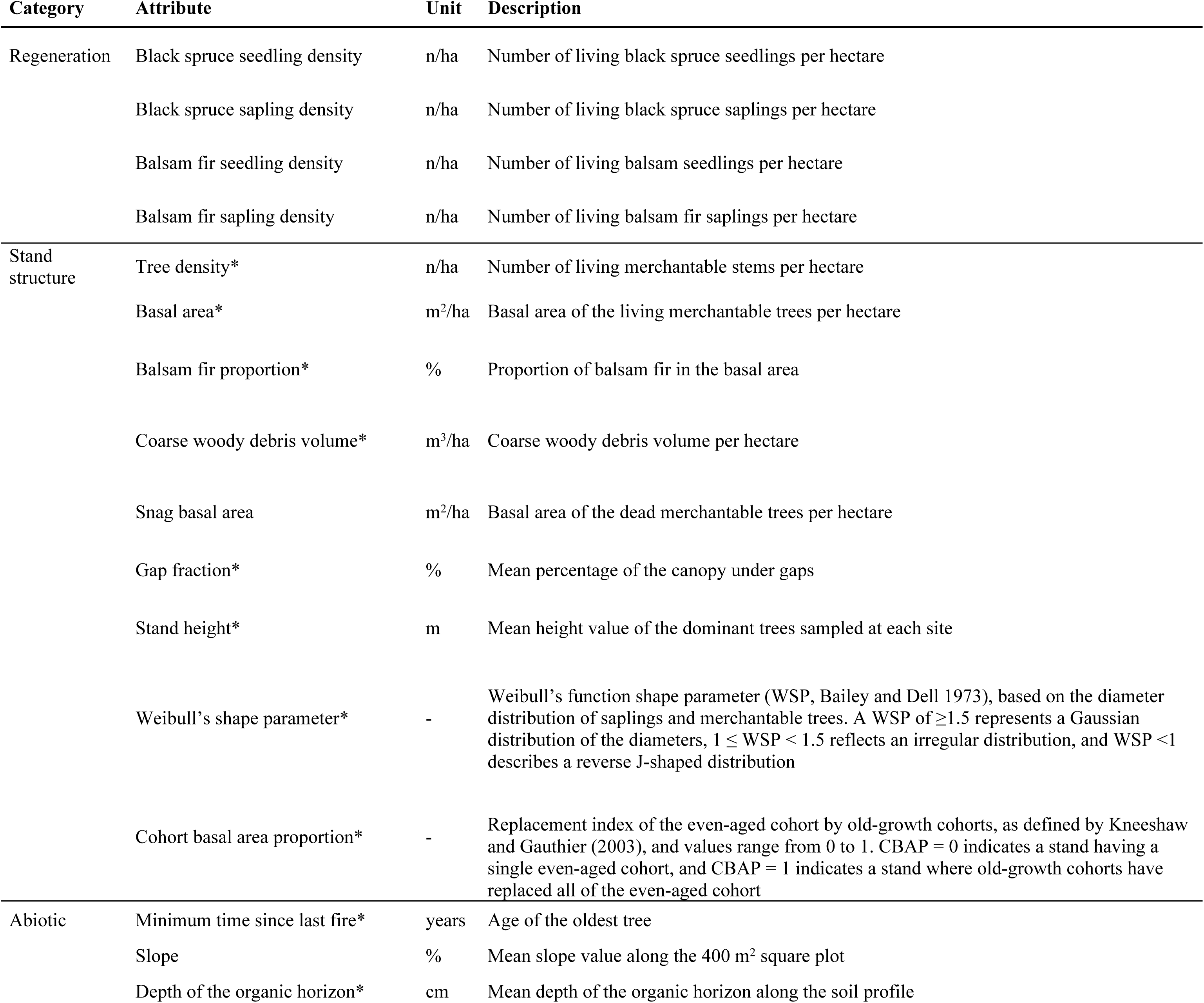
Description of the regeneration, stand structure, and abiotic attributes sampled at the study sites as adapted from Martin et al. (2018). “*” indicates attributes computed by Martin et al. (2018).

### Data analysis

First, we performed k-means clustering (53) on black spruce and balsam fir regeneration attributes to identify the main patterns driving the regeneration dynamics of these two tree species in eastern Canadian boreal old-growth forests. To highlight the differences between the two species, we ran k-means clustering for each species separately. The clustering of black spruce regeneration relied on black spruce seedling and sapling densities of all 71 sites. Similarly, clustering of balsam fir regeneration also relied on balsam seedling and sapling densities; however, balsam fir seedlings and saplings were absent in 24 sites. We thus removed these sites for the clustering of balsam fir (47 plots remaining) to eliminate any influence from sites lacking this balsam fir regeneration. For each cluster analysis, we determined the optimal number of regeneration clusters using the simple structure index (SSI; 54) criterion. Separately for both species, we compared regeneration as well as the structural and environmental attributes within the clusters. We used analysis of variance (ANOVA) when the ANOVA conditions were fulfilled (data normality and homoscedasticity) or Kruskall-Wallis nonparametric analysis of variance when these conditions were not met. When ANOVA or the Kruskall-Wallis tests were significant, we performed a Tukey posthoc test (55) or a Fisher’s least significant difference test (56), respectively. Moreover, we also calculated Spearman’s rank correlation coefficient between the regeneration and structural/environmental attributes. This latter analysis aimed to provide valuable information for interpreting our results by highlighting the strength of the relationship between regeneration and these various attributes.

All analyses were performed using R software, version 3.3.1 (57) and the *vegan* (58), *Hmisc* (59), and *agricolae* (60) packages, applying a *p*-threshold of 0.05.

## Results

### Black spruce and balsam fir regeneration

For cluster analysis of black spruce regeneration, we determined eight as being the optimal number of clusters (SSI = 2.23; **Fig 3**). Black spruce seedling and sapling densities differed significantly between the black spruce regeneration clusters (BS; **Table 2A**). Black spruce seedling density was more than 8× higher in cluster BS8, having the highest density (26 543 seedlings/ha), than in cluster BS1, characterized by the lowest seedling density values (3 008 seedlings/ha). Black spruce seedling density did not differ between clusters BS4, BS5, and BS6. Regarding the density of black spruce saplings, cluster BS1—having the lowest values at 322 saplings/ha—contained a sapling density 12× less than cluster BS5, which had the highest density of black spruce seedlings at 3 783 saplings/ha. The remaining clusters, characterized by intermediate values of black spruce sapling density, aligned along a gradient. We also observed significant differences in balsam fir seedling density between clusters. Concerning balsam fir seedling density within the clusters of black spruce regeneration, we observed significant differences, ranging from 873 seedlings/ha (lowest value, cluster BS7) to 9 720 seedlings/ha (highest value, cluster BS1); however, balsam fir sapling density did not differ significantly between the clusters.

**Fig 3.**
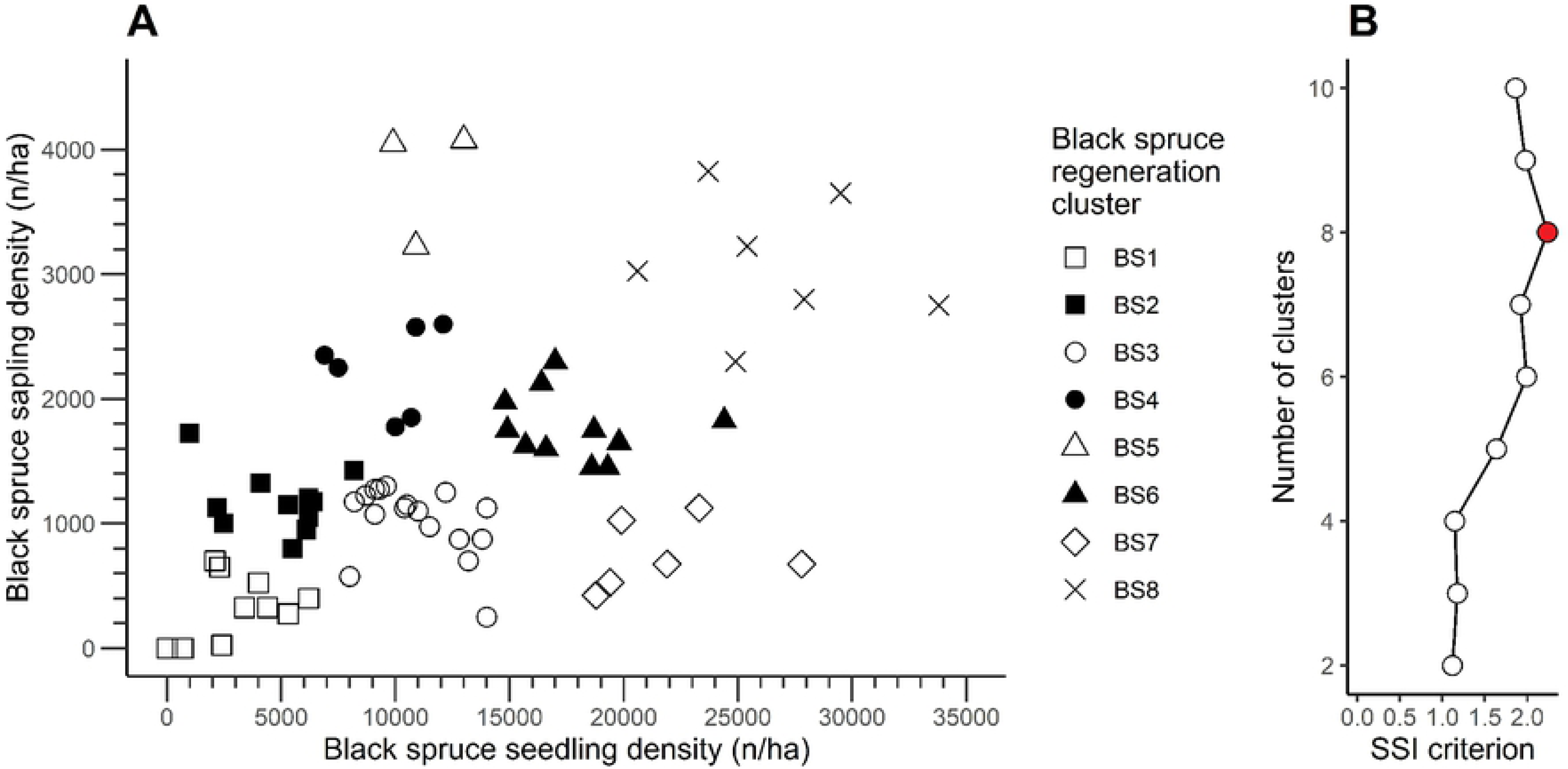
(A) Density of black spruce seedlings and saplings at the 71 studied sites, grouped by black spruce regeneration clusters. (B) Value of the SSI criterion according to the number of clusters for black spruce using k-means clustering. Filled circle in (B) indicates the highest value of the SSI criterion.

**Table 2.** Mean and standard error of the regeneration attributes for (A) black spruce regeneration clusters and (B) balsam fir regeneration clusters. Different letters indicate significant differences at *p* < 0.05, following a > b > c > d > e. BS: black spruce; BF: Balsam fir

**A: Black spruce regeneration**
| Cluster | BS1 (n=10) | BS2 (n = 11) | BS3 (n = 17) | BS4 (n = 6) | BS5 (n = 3) | BS6 (n = 11) | BS7 (n = 6) | BS8 (n = 7) |
| --- | --- | --- | --- | --- | --- | --- | --- | --- |
| Black spruce seedling density (n/ha) | 3 080 ± 1 959 e | 4 882 ± 2 177 d | 10 906 ± 2 096 c | 9 683 ± 2 048 c | 11 267 ± 1 582 c | 17 836 ± 2 777 b | 21 850 ± 3 365 a | 26 543 ± 4 295 a |
| Black spruce sapling density (n/ha) | 322 ± 257 f | 1 175 ± 251 d | 1 019 ± 286 d | 2 233 ± 353 bc | 3 783 ± 484 a | 1 773 ± 269 c | 742 ± 277 e | 3 082 ± 532 ab |
| Balsam fir seedling density (n/ha) | 9 720 ± 7 920 a | 5 773 ± 7 621 ab | 1 835 ± 4 253 c | 1 917 ± 4 224 c | 3 333 ± 5 687 ac | 873 ± 2 039 c | 6 317 ± 8 655 ab | 2 357 ± 5 474 bc |
| Balsam fir sapling density (n/ha) | 1 492 ± 1 499 | 2 516 ± 3 374 | 228 ± 495 | 500 ± 765 | 608 ± 1 032 | 125 ± 357 | 592 ± 668 | 200 ± 416 |

**B: Balsam fir regeneration**
| Cluster | BF1 (n = 28) | BF2 (n = 11) | BF3 (n = 3) | BF4 (n = 5) |
| --- | --- | --- | --- | --- |
| Black spruce seedling density (n/ha) | 14 379 ± 7 824 a | 10 773 ± 9 688 ab | 1 900 ± 794 b | 9 180 ± 7 258 ab |
| Black spruce sapling density (n/ha) | 1 670 ± 1 025 a | 1 282 ± 1 131 ab | 1 283 ± 388 ab | 470 ± 151 b |
| Balsam fir seedling density (n/ha) | 957 ± 1 520 c | 9 827 ± 3 243 b | 1 6267 ± 5 460 ab | 1 8740 ± 2 756 a |
| Balsam fir sapling density (n/ha) | 223 ± 397 c | 1 464 ± 683 b | 7 442 ± 1 934 a | 2 590 ± 1 066 ab |

Regarding balsam fir regeneration, two and four clusters produced an identical SSI criterion value of 1.14; **Fig 4**. Nonetheless, to obtain a more detailed evaluation of the dynamics of balsam fir regeneration, we chose to use four clusters (BF; **Table 2B**). Balsam fir seedling and sapling density varied markedly between clusters, and we identified significant differences for every attribute between the clusters. For example, the density of balsam fir seedlings within cluster BF4, marked by the highest seedling density at 18 740 seedlings/ha, was almost 20× that of the cluster having the lowest density of balsam fir seedlings (957 seedlings/ha; cluster BF1).

**Fig 4.**
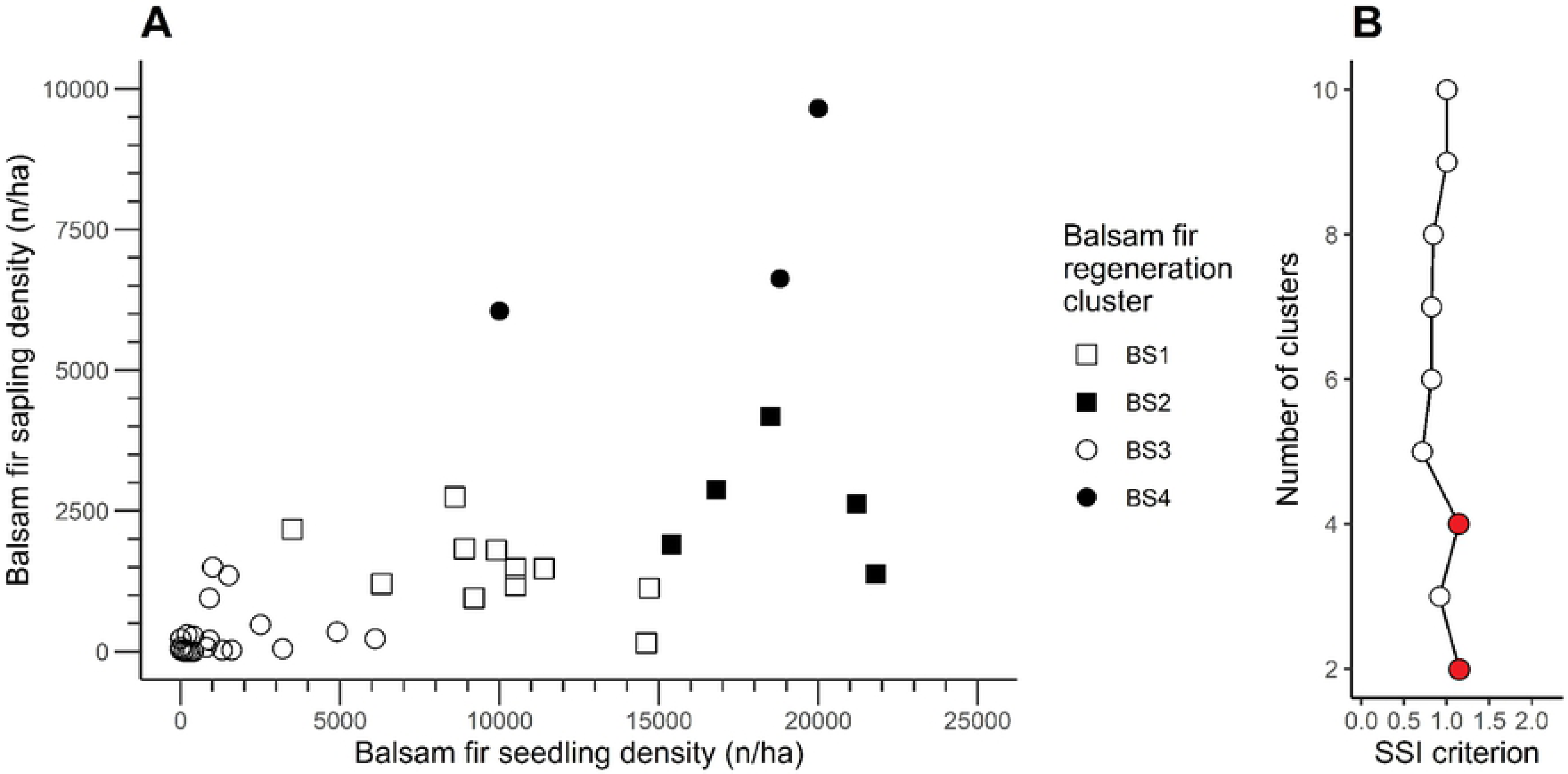
(A) Density of balsam fir seedlings and saplings at the 48 studied sites of the balsam fir regeneration portion of the study, grouped by balsam fir regeneration clusters. (B) Value of the SSI criterion according to the number of clusters for balsam fir using k-means clustering. Filled red circles in (B) indicate the highest value of the SSI criterion.

Similarly, the highest density of balsam fir saplings (7 442 saplings/ha; cluster BF3) was 33× that of the cluster having the lowest density (223 saplings/ha; cluster BF1). Differences between clusters in terms of black spruce seedling or sapling density were less striking, although both attributes differed significantly between the clusters. Black spruce seedling density varied from 1 900 to 14 379 seedlings/ha, whereas black spruce sapling density ranged from 470 to 1 670 saplings/ha (clusters BF4 and BF1, respectively, for both cases).

### Structural and environmental attributes

Densities of black spruce seedlings and saplings both correlated positively with gap fraction, cohort basal area proportion, minimum time since the last fire, and depth of the organic horizon; both correlated negatively with slope (**Table 3**). Black spruce seedling density correlated negatively with basal area, balsam fir proportion, and maximum height. Balsam fir seedling and sapling densities correlated positively with balsam fir proportion, coarse woody debris volume, snag basal area, maximum height, and slope. Balsam fir seedling density also correlated significantly with basal area. In general, correlation coefficients tended to be relatively low even when significant; this was especially true for black spruce as no correlation coefficient between sapling density and gap fraction exceeded 0.5. These relatively low coefficient values indicate a relatively weak relationship between black spruce regeneration and the structural and environmental attributes. We observed, however, elevated correlation coefficients (≥0.5) for balsam fir in relation to several structural and environmental attributes, including balsam fir proportion, slope, coarse woody debris volume (saplings only), and snag basal area (seedlings only).

**Table 3.** Spearman correlation coefficients between regeneration attributes and structural and environmental attributes. “*” indicates significance at *p* < 0.05, “**” at *p* < 0.01 and “***” at *p* < 0.001.

| Category | Attribute | Black spruce |  | Balsam fir |  |
| --- | --- | --- | --- | --- | --- |
|  |  | Seedlings | Saplings | Seedlings | Saplings |
| Structure | Tree density (n/ha) | 0.18 | -0.09 | 0.14 | 0.10 |
|  | Basal area (m<sup>2</sup>/ha) | -0.09 | -0.49*** | 0.36* | 0.17 |
|  | Balsam fir proportion (%) | -0.21 | -0.26* | 0.80*** | 0.86*** |
|  | Gap fraction (%) | 0.34** | 0.51*** | -0.17 | -0.02 |
|  | Weibull's shape parameter | 0.05 | 0.16 | -0.19 | -0.09 |
|  | Coarse woody debris volume (m<sup>3</sup>/ha) | -0.11 | -0.07 | 0.44** | 0.61*** |
|  | Snag basal area (m<sup>2</sup>/ha) | -0.21 | -0.21 | 0.55*** | 0.48*** |
|  | Maximum height (m) | -0.12 | -0.31** | 0.39** | 0.33* |
|  | Cohort basal area proportion | 0.32** | 0.24* | 0.08 | 0.07 |
| Abiotic | Minimum time since the last fire (years) | 0.43*** | 0.32** | -0.19 | -0.22 |
|  | Slope (%) | -0.29* | -0.37** | 0.59*** | 0.56*** |
|  | Depth of the organic horizon (cm) | 0.41*** | 0.32** | -0.17 | -0.10 |

Black spruce regeneration clusters differed significantly from each other for many attributes, including basal area, gap fraction, minimum time since the last fire, slope, and depth of the organic horizon (**Table 4**). We identified marked differences between the study attributes and clusters; for example, basal area differed two-fold between cluster BS8 and cluster BS7, gap fraction values of cluster BS1 were more than double those of cluster BS5, the minimum time since the last fire varied from 146 (cluster BS1) to 249 years (cluster BS8), cluster BS8 has a 5× higher slope than that of cluster BS1 (4.0% versus 23.4%, respectively), organic horizon depth varied from 16.0 cm (cluster BS1) to 47.9 cm (cluster BS8). Overall, clusters BS1 and BS8 were the most distinct clusters; the other clusters fell along a gradient between this pair of clusters. Cluster BS1 grouped stands located on steeper sites, characterized by a shallow organic horizon, a dense canopy, a high basal area, and relatively young trees. In contrast, cluster BS8 grouped stands having a gentle slope as well as a thick organic horizon, open canopy, low basal area, and older trees. The remaining clusters represented intermediate values between these two boundary clusters.

**Table 4.**
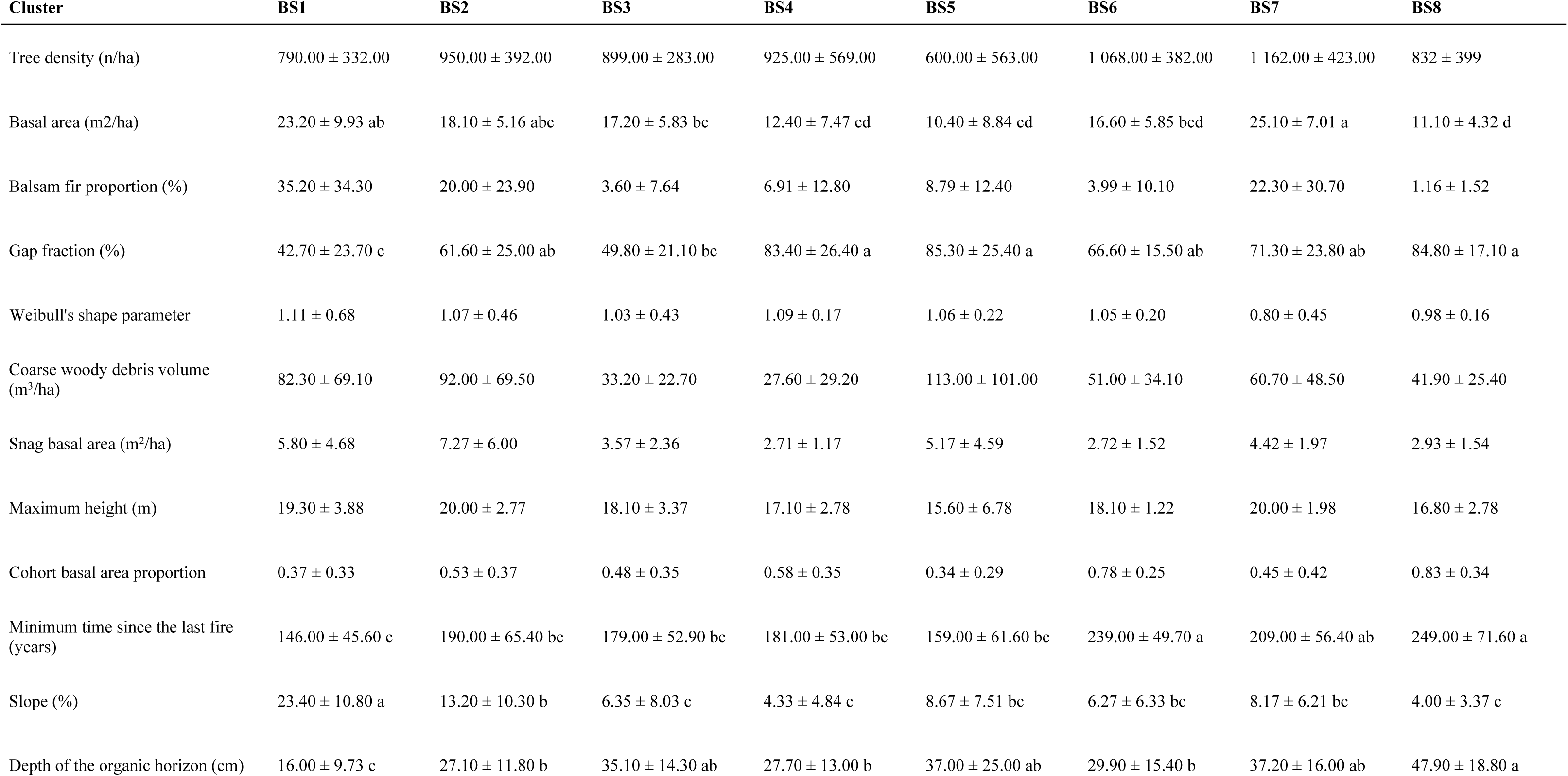
Mean and standard error of the structural and environmental attributes for black spruce regeneration clusters (BS). Different letters indicate significant differences at *p* < 0.05, following a > b > c > d. BS: black spruce; BF: Balsam fir

| Cluster | BS1 | BS2 | BS3 | BS4 | BS5 | BS6 | BS7 | BS8 |
| --- | --- | --- | --- | --- | --- | --- | --- | --- |
| Tree density (n/ha) | 790.00 ± 332.00 | 950.00 ± 392.00 | 899.00 ± 283.00 | 925.00 ± 569.00 | 600.00 ± 563.00 | 1 068.00 ± 382.00 | 1 162.00 ± 423.00 | 832 ± 399 |
| Basal area (m<sup>2</sup>/ha) | 23.20 ± 9.93 ab | 18.10 ± 5.16 abc | 17.20 ± 5.83 bc | 12.40 ± 7.47 cd | 10.40 ± 8.84 cd | 16.60 ± 5.85 bcd | 25.10 ± 7.01 a | 11.10 ± 4.32 d |
| Balsam fir proportion (%) | 35.20 ± 34.30 | 20.00 ± 23.90 | 3.60 ± 7.64 | 6.91 ± 12.80 | 8.79 ± 12.40 | 3.99 ± 10.10 | 22.30 ± 30.70 | 1.16 ± 1.52 |
| Gap fraction (%) | 42.70 ± 23.70 c | 61.60 ± 25.00 ab | 49.80 ± 21.10 bc | 83.40 ± 26.40 a | 85.30 ± 25.40 a | 66.60 ± 15.50 ab | 71.30 ± 23.80 ab | 84.80 ± 17.10 a |
| Weibull's shape parameter | 1.11 ± 0.68 | 1.07 ± 0.46 | 1.03 ± 0.43 | 1.09 ± 0.17 | 1.06 ± 0.22 | 1.05 ± 0.20 | 0.80 ± 0.45 | 0.98 ± 0.16 |
| Coarse woody debris volume (m<sup>3</sup>/ha) | 82.30 ± 69.10 | 92.00 ± 69.50 | 33.20 ± 22.70 | 27.60 ± 29.20 | 113.00 ± 101.00 | 51.00 ± 34.10 | 60.70 ± 48.50 | 41.90 ± 25.40 |
| Snag basal area (m<sup>2</sup>/ha) | 5.80 ± 4.68 | 7.27 ± 6.00 | 3.57 ± 2.36 | 2.71 ± 1.17 | 5.17 ± 4.59 | 2.72 ± 1.52 | 4.42 ± 1.97 | 2.93 ± 1.54 |
| Maximum height (m) | 19.30 ± 3.88 | 20.00 ± 2.77 | 18.10 ± 3.37 | 17.10 ± 2.78 | 15.60 ± 6.78 | 18.10 ± 1.22 | 20.00 ± 1.98 | 16.80 ± 2.78 |
| Cohort basal area proportion | 0.37 ± 0.33 | 0.53 ± 0.37 | 0.48 ± 0.35 | 0.58 ± 0.35 | 0.34 ± 0.29 | 0.78 ± 0.25 | 0.45 ± 0.42 | 0.83 ± 0.34 |
| Minimum time since the last fire (years) | 146.00 ± 45.60 c | 190.00 ± 65.40 bc | 179.00 ± 52.90 bc | 181.00 ± 53.00 bc | 159.00 ± 61.60 bc | 239.00 ± 49.70 a | 209.00 ± 56.40 ab | 249.00 ± 71.60 a |
| Slope (%) | 23.40 ± 10.80 a | 13.20 ± 10.30 b | 6.35 ± 8.03 c | 4.33 ± 4.84 c | 8.67 ± 7.51 bc | 6.27 ± 6.33 bc | 8.17 ± 6.21 bc | 4.00 ± 3.37 c |
| Depth of the organic horizon (cm) | 16.00 ± 9.73 c | 27.10 ± 11.80 b | 35.10 ± 14.30 ab | 27.70 ± 13.00 b | 37.00 ± 25.00 ab | 29.90 ± 15.40 b | 37.20 ± 16.00 ab | 47.90 ± 18.80 a |

We noted significant differences between balsam fir regeneration clusters in terms of balsam fir proportion, coarse woody debris volume, snag basal area, and slope (**Table 5**). As with the black spruce regeneration clusters, two balsam fir regeneration clusters—clusters BF1 and BF4— represented opposite extremes along a gradient. Balsam fir proportion was almost 14× higher in cluster BF4 (56.7%) than in cluster BF1 (4.12%). Coarse woody debris volume in cluster BF3 was more than double that of cluster BF1, at 61.6 and 155 m3/ha, respectively. Cluster BF4 contained a snag basal area that was more than triple that of cluster BF1 (14 versus 3.9 m^2^/ha, respectively). Slope in cluster BF4 (28.4%) was also 4× higher than that in cluster BF1 (8.14%). All told, cluster BF1 represented sites having a gentle slope and lower balsam fir proportion, as well as a moderate coarse woody debris volume and snag basal area. Cluster BF3, on the other hand, grouped sites marked by steeper slopes, as well as higher values of balsam fir proportion, coarse woody debris volume, and snag basal area. As above, the remaining clusters fell between these two extreme clusters. Relative to the black spruce results, however, these two balsam fir clusters differed much less from each other; for example, we observed no significant differences in coarse woody debris volume for clusters BF2, BF3, and BF4. This pattern implies that the structural differences within the balsam fir regeneration clusters were less noticeable than those observed in the black spruce stands.

**Table 5.** Mean ± standard deviation of structural and environmental attributes for balsam fir regeneration clusters (BF). Letters indicate significant differences at *p* < 0.05, following a > b > c.

| Cluster | BF1 | BF2 | BF3 | BF4 |
| --- | --- | --- | --- | --- |
| Tree density (n/ha) | 880.00 ± 332.00 | 927.00 ± 277.00 | 892.00 ± 104.00 | 810.00 ± 326.00 |
| Basal area (m<sup>2</sup>/ha) | 17.50 ± 7.27 | 20.80 ± 7.64 | 14.90 ± 1.92 | 21.70 ± 6.94 |
| Balsam fir proportion (%) | 4.12 ± 6.95 b | 28.80 ± 19.10 a | 55.00 ± 5.10 a | 56.70 ± 35.40 a |
| Gap fraction (%) | 64.10 ± 26.00 | 57.10 ± 29.80 | 72.70 ± 14.80 | 64.00 ± 27.10 |
| Weibull's shape parameter | 0.87 ± 0.29 | 1.15 ± 0.62 | 0.88 ± 0.12 | 0.81 ± 0.13 |
| Coarse woody debris volume (m<sup>3</sup>/ha) | 61.60 ± 47.00 b | 84.00 ± 35.60 a | 155.00 ± 62.90 a | 121.00 ± 60.00 a |
| Snag basal area (m<sup>2</sup>/ha) | 3.90 ± 3.05 c | 5.09 ± 2.00 bc | 14.00 ± 5.58 a | 7.97 ± 4.40 ab |
| Maximum height (m) | 18.90 ± 3.05 | 20.70 ± 2.06 | 19.70 ± 2.23 | 21.20 ± 1.75 |
| Cohort basal area proportion | 0.60 ± 0.35 | 0.63 ± 0.36 | 0.82 ± 0.30 | 0.74 ± 0.16 |
| Minimum time since the last fire (years) | 213.00 ± 66.50 | 193.00 ± 50.60 | 188.00 ± 50.10 | 204.00 ± 41.40 |
| Slope (%) | 8.14 ± 9.11 c | 12.50 ± 10.5 bc | 18.70 ± 5.03 ab | 28.40 ± 6.02 a |
| Depth of the organic horizon (cm) | 31.60 ± 16.00 | 26.10 ± 14.00 | 29.00 ± 15.10 | 21.60 ± 9.29 |

## Discussion

Old-growth forests are critical habitats for biodiversity and ecosystem services. A better understanding of their functioning is therefore necessary for developing sustainable management strategies. The results of our study highlight that regeneration in boreal old-growth forests involves complex processes (non-linear, self-organized, disturbance-driven, structurally-dependent, etc.) that cannot be summarized along a single linear chronosequence of forest succession or by using a limited number of structural attributes as proxies. In general, we observed secondary disturbance regimes and topographic constraints as the main drivers of balsam fir and black spruce regeneration in our study stands. Temporal and spatial scales are therefore two important factors to explain the dynamics of tree regeneration in the boreal old-growth forests of eastern Canada.

### Dynamics of black spruce regeneration

The dynamics of black spruce regeneration in boreal old-growth forests involve highly complex processes. We observed highly variable seedling and sapling densities within the study stands, and specific structural attributes defined each black spruce regeneration cluster. These observations may explain the low Spearman correlation coefficients observed for black spruce, as its regeneration density depends on multiple and interrelated factors (10,15,61). Moreover, the black spruce regeneration clusters present no significant differences in their cohort basal area proportions; therefore, differences between clusters did not result from succession toward an old-growth stage. We observed a significant difference between clusters in relation to minimum time since the last fire; however, this value generally exceeded 150 years, i.e., the threshold beyond which tree age becomes a poor indicator of stand age in boreal forests (62, 63). As such, changes in stand structure due to secondary disturbance are more relevant for explaining regeneration dynamics rather than invoking the process of forest succession.

For black spruce, differences in the structural attributes between regeneration clusters testify to the influence of disturbance on seedling and sapling density (**Fig 5A**). As a starting point, cluster BS7 grouped dense old-growth forest stands found on gentle to medium slopes (0–7% and 8– 24%, respectively). The stands in this cluster contained a moderate gap fraction and a high basal area, i.e., stands that have neither been recently nor significantly disturbed. Indeed, due to their narrow canopy, even dense old-growth black spruce stands can be characterized by a relatively high gap fraction (41). At this cluster’s successional stage, a low black spruce sapling density and high seedling density indicated a dense understory waiting for a canopy opening. This distribution of trees, saplings, and seedlings agrees with previous results (41, 63) that identified a low suppressed tree density in old-growth stands that had a dense canopy and that were dominated by black spruce. It is quite likely that most of the black spruce seedlings sampled in the study sites represented layers rather than seeds. Indeed, this regeneration strategy is more effective on soils where most of the organic horizon is covered by a layer of mosses and organic matter (65). Moreover, these layers generally remain connected to the mother tree at this seedling stage and, thus, these layers likely remain under hormonal control with the process of apical dominance inhibiting their growth (lateral growth) (66–68).

**Fig 5.**
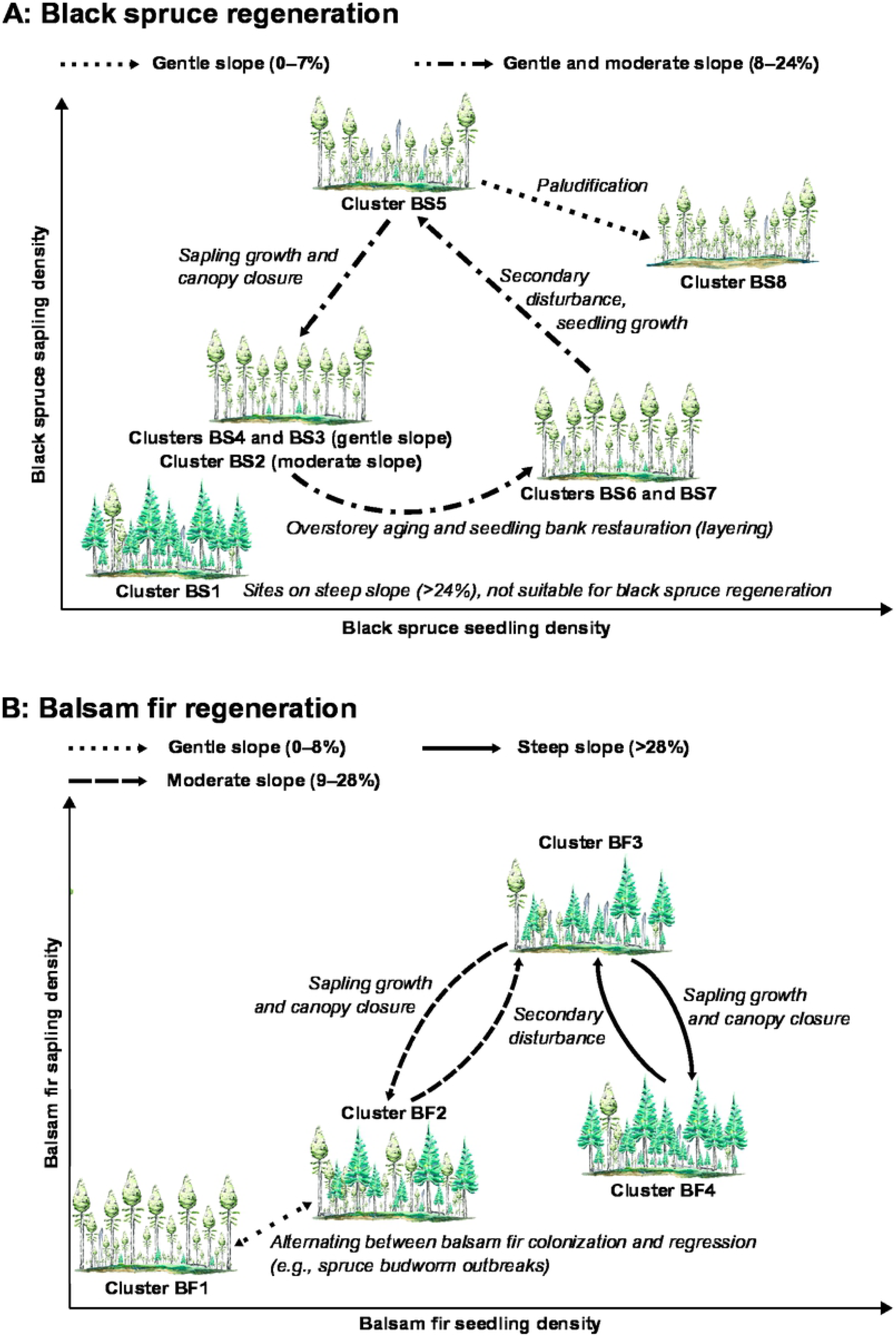
Dynamics of (A) black spruce and (B) balsam fir regeneration according to secondary disturbance regime and topography as derived from the identified regeneration clusters. Water paintings by Valentina Buttò.

Overstory trees aged progressively and became increasingly sensitive to secondary disturbances and senescence-induced mortality (69–71). Cluster BS7 became cluster BS6, and black spruce sapling density began to increase. Overstory trees eventually died, creating gaps and decreasing the stand basal area. Black spruce regeneration individuals, including layers, are efficient gap-fillers (26,30,33), and these layers are no longer subject to apical control upon the death of the mother tree. Hence, most seedlings benefited from these openings to produce to a high sapling density, i.e., cluster BS6 shifted to cluster BS5. Saplings eventually reached the overstory and progressively closed the canopy. The result was a significant decrease in sapling density.

However, we observed two different pathways depending on stand topography: gentle slopes (clusters BS4 and BS3, sapling growth and canopy closure, respectively) and moderate slopes (cluster BS2, sapling growth and canopy closure). Canopy closing finally led to an increased stand basal area, i.e., clusters BS2 and BS3 shifted toward cluster BS7, reinitiating the cycle. While we observed few changes in black spruce sapling density during this last transition, seedling density increased sharply, indicating the re-establishment of a dense understory layer awaiting the next canopy opening.

The two remaining black spruce regeneration clusters both represented two specific abiotic conditions and dynamics. BS8 was defined by a gentle slope, a thick organic horizon, a high gap fraction, and a low basal area. These characteristics typify stands undergoing paludification—the accumulation of soil organic matter due to insufficient drainage resulting in a decreased stand productivity (72, 73). Paludification inhibits tree growth, but not black spruce regeneration. As a result, black spruce sapling and seedling densities are often dense in paludified black spruce stands, but these saplings and seedlings are unable to close the gaps caused by overstory tree death (29). Paludification, however, is a process limited to specific conditions, i.e., poor drainage and low temperatures; this process is not observed within well or moderately well-drained soils, i.e., stands having a minimum slope (74–76), explaining, therefore, the particularity of this cluster.

BS1, on the other hand, was defined by a shallow organic horizon, a steep slope, and a low gap fraction. This cluster presented the lowest black spruce seedling and sapling densities; this pattern matches prior observations in the study area that the abundance of black spruce decreases progressively as slope increases, eventually being replaced by balsam fir and northern hardwoods (40, 45). Competition with balsam fir could explain the limited regeneration of black spruce on these steepest sites. However, another factor could be the thin organic horizon that reduces the survival of black spruce layers due to insufficient moisture, especially in the summer (77).

Nonetheless, in sufficiently drained sites of more moderate slope, black spruce regeneration in old-growth forests presented a dynamic having four phases: 1) development of a dense seedling bank under a closed canopy; 2) rapid seedling growth once the overstory was disturbed and causing a decrease in seedling density and an increase in sapling density; 3) progressive canopy closure, implying a decrease in sapling density as saplings become merchantable trees; and 4) a return to phase 1.

### Balsam fir regeneration dynamics

Disentangling balsam fir regeneration dynamics in the study stands presented a greater challenge than that for black spruce dynamics as balsam fir regeneration was absent for 24 plots and sparse for the 28 sites belonging to cluster BF1. Several factors may explain the scarcity of balsam fir regeneration in most of the studied stands, such as soils being too wet or the stands having a limited seed bank. In the sites characterized by relatively poor drainage, very wet and cold soils inhibit balsam fir seed germination and favor black spruce layering (78, 79). In the study region, the fire cycle is shorter in the valley bottoms than on the hilltops (45), probably due to a later snowmelt at higher elevations. Balsam fir is not a fire-adapted species, and this tree often requires decades if not centuries to recolonize a burned area (80). Moreover, the dispersal of balsam fir seeds is relatively limited, and its occurrence requires proximal seed trees (15, 39) as evidenced by the strong correlation observed between the proportion of balsam fir and the balsam fir regeneration density. Shorter fire cycles in the valley bottoms may thus inhibit the colonization of balsam fir in these areas of the study territory. Nevertheless, the absence of balsam fir in boreal old-growth stands is common in eastern Canada (19,20,40) because of all the factors explained previously; as such, sampling bias does not account for the results in our study.

We observed no significant difference between the balsam fir regeneration clusters in terms of the minimum time since the last fire and the cohort basal area proportion. As with the black spruce clusters, all balsam fir clusters represented the old-growth successional stage. Previous research of balsam fir regeneration dynamics in the boreal forests of eastern Canada focused on stands at the beginning of the transition toward the old-growth stage (e.g., 27,80,81). Our results underscore that once the old-growth stage is attained, and if seed trees are present nearby, the existing seed bank is sufficient to provide continuous regeneration of balsam fir (28, 83).

Moreover, we observed significantly different stand slopes between the clusters, highlighting the importance of topography in explaining balsam fir stand dynamics (40). These results imply that as in the case of black spruce, secondary disturbance dynamics and topographic constraints drive balsam fir regeneration in the old-growth forests of eastern Canada.

For sites located on gentle slopes (0–8%), we observed two different balsam fir regeneration clusters. One cluster represented sites where balsam fir was almost absent from the canopy (BF1), whereas the other cluster represented stands where balsam fir accounted for around 30% of the basal area (BF2). As a result, there was almost no balsam fir regeneration in BF1, while seedling and sapling densities were of moderate levels in BF2. Coarse woody debris volume was, however, higher in BF2 than BF1, suggesting more recent disturbances (**Fig 5B**). This involves a dynamic where boreal old-growth species composition switches between a pure black spruce stand and a mixed black spruce and balsam fir stand, possibly with the presence of white birch at a very low abundance (27, 28). This type of dynamic is consistent with previous observations (28, 40). Balsam fir is a competitive species that can quickly reach the upper canopy following a secondary disturbance (28, 31). It is also very sensitive to disturbance, especially spruce budworm outbreaks, the main secondary disturbance agent in eastern Canadian boreal forests (37,84,85). Outbreaks of this insect heighten balsam fir mortality as spruce budworm larvae emergence is well synchronized with balsam fir budburst. In contrast, black spruce mortality during spruce budworm outbreaks is relatively low as black spruce budburst and larval emergence are poorly synchronized (86). The most severe budworm outbreaks cause significant mortality of the regeneration, in particular that of balsam fir (38,87,88). As a result, balsam fir abundance may decrease significantly in formerly mixed black spruce–balsam fir stands following an outbreak, although balsam fir may, with time, progressively recolonize the stand (20, 21).

We observed no difference between the balsam fir regeneration clusters BF2 and BF3 in terms of coarse woody debris volume and proportion of balsam fir; this pattern represents dynamics in sites of moderate slope (i.e., 9–28%). However, the snag basal area was significantly higher in BF3. Relative to black spruce, balsam fir is also more vulnerable to windthrow and fungal rot (35, 36). The presence of an important coarse woody debris volume in stands with an elevated balsam fir proportion in the canopy is therefore consistent with balsam fir ecology. However, a higher snag basal area can also indicate a relatively recent disturbance, as black spruce and balsam fir snags often fell in the twenty years following a tree death (89). Hence, cluster BF3 may group recently disturbed stands marked by a dynamic balsam fir regeneration that quickly fills the canopy (27,28,31). Once the canopy is closed, stand structure shifts to BF2, defined by a dense seedling bank.

Finally, BF3 and BF4 grouped stands on steep slopes (>28%), yet that no had significant structural differences between the two clusters. This result may reflect the low number of sites sampled for both clusters (3 and 5 sites, respectively). However, it is also probable that they represented a balsam fir regeneration dynamic similar to that observed on moderate slopes, with BF3 grouping recently disturbed stands and the BF4 grouping the resilient stands. On intermediate slopes, black spruce regeneration continued to compete with balsam fir, thereby explaining the intermediate balsam fir seedling density in BF2. On steep slopes, however, balsam fir dominated the canopy. It is therefore likely that these stands were driven by regular small- and moderate-scale disturbances (26), resulting in recurrent deadwood inputs and active regeneration/mortality phases.

## Conclusion

This study determined how secondary disturbance regimes and topographic constraints explain the dynamics of black spruce and balsam fir regeneration in old-growth forests. Thus, our study refutes a classic assumption in forest science by demonstrating that the standard linear and theoretical paradigms (successional stages) are not able to explain the complexity of old-growth forest dynamics where many ecological factors interact at multiple temporal and spatial scales. Second, this study provides a better acknowledgment of the importance of regeneration dynamics in the boreal old-growth forests of eastern Canada. Disturbance dynamics in these ecosystems are, however, defined by disturbances that vary in terms of type, frequency, and severity (26, 71). Thus, our results highlight the overall trends of regeneration dynamics in old-growth forests, and further research is required to determine how these trends may change depending on disturbance characteristics.

Third, sustainable forest management aims to develop new silvicultural treatments to minimize differences between natural and managed stands. For this, partial cuttings offer a promising solution to adapt forestry practices to act in a similar manner as secondary disturbance regimes. These treatments, however, must be adapted to conditions within the eastern Canadian forest (90–93). The results of our study provide new guidelines for a forest management approach that brings the regeneration dynamics within managed stands closer to those of boreal old-growth forests.

## Acknowledgments

We thank Audrey Bédard, Jean-Guy Girard, Émilie Chouinard, Anne-Élizabeth Harvey, Aurélie Cuvelière, Évelyn Beliën, and Angelo Fierravanti for their precious help during field sampling. Yan Boucher and Pierre Grondin from Quebec’s Ministry of Forests, Fauna, and Parks (MFFP) provided the data collected from the study territory. We also thank Valentina Buttò for her water paintings, used in Figure 4 of this paper.

